# Crystal structure of the giant panda MHC class I complex: first insights into the viral peptide presentation profile in the bear family

**DOI:** 10.1101/2020.01.15.908608

**Authors:** Hongyu Yuan, Lizhen Ma, Lijie Zhang, Xiaoying Li, Chun Xia

**Affiliations:** Department of Microbiology and Immunology, College of Veterinary Medicine, China Agricultural University, Haidian District, Beijing 100193, China; National Laboratory of Biomacromolecules, Institute of Biophysics, Chinese Academy of Sciences, Beijing 100101, China; School of Basic Medical Sciences, Xinxiang Medical University, Xinxiang, Henan, China

**Keywords:** Crystal structure, Giant panda, MHC-I, Peptide presentation, Evolution

## Abstract

The viral cytotoxic T lymphocyte (CTL) epitope peptides presented by classical MHC-I molecules require the assembly of a peptide-MHC-I-*β*2m (aka pMHC-I) trimolecular complex for TCR recognition, which is the critical activation link for triggering antiviral T cell immunity. Ursidae includes 5 genera and 8 species; however, research on T cell immunology in this family, especially structural immunology, is lacking. In this study, the structure of the key trimolecular complex pMHC-1 (aka pAime-128), which binds a peptide from canine distemper virus, was solved for the first time using giant panda as a representative species of Ursidae. The structural characteristics of the giant panda pMHC-I complex, including the unique pockets in the peptide-binding groove (PBG), were analyzed in detail. Comparing the panda pMHC-I to others in the bear family and extending the comparison to other mammals revealed distinct features. The interaction between MHC-I and *β*2m, the features of pAime-128 involved in TCR docking and CD8 binding, the anchor sites in the PBG, and the CTL epitopes of potential viruses that infect pandas were concretely clarified. Unique features of pMHC-I viral antigen presentation in the panda were revealed by solving the three-dimensional structure of pAime-128. The distinct characteristics of pAime-128 indicate an unusual event that emerged during the evolution of the MHC system in the bear family. These results provide a new platform for research on panda CTL immunity and the design of vaccines for application in the bear family.

**IMPORTANCE:** Ursidae includes 5 genera and 8 species; however, the study of its immunology, especially structural immunology, is extremely rare to date. In this paper, we first crystallized the key complex pMHC-I, taking the giant panda as its representative species. Structural characteristics of the giant panda pMHC-I complexes, contains the unique pockets of PBG were analyzed in detail. Comparison of the panda pMHC-I in the bear family and other mammals, almost definite features was displayed. Meanwhile, the interaction between HC and LV, the unique features of pMHC-I in the CD8 binding and TCR docking, validation of anchor site in the PBG, and epitopes of potential viruses infected with the pandas, were concretely clarified. These unique characteristics of pMHC-I clearly indicate an unusual situation during the evolution of MHC molecules in the endangered pandas. These results also provide a novel platform for further study of panda T cell immunology and vaccines.

## INTRODUCTION

The cytotoxic T lymphocytes (CTLs) and their immune responses are common to various genera of jawed vertebrates. Common CTLs include CD8^+^ T cells, which recognize only antigens presented by classical major histocompatibility complex class I (MHC-I) molecules (1). MHC-I genes are expressed in most nucleated cells and present antigens derived mostly from intracellular viruses, parasites, and tumors (2). Structural studies of MHC-I molecules have shown that the complex is generally composed of an MHC-I, *β*2-microglobulin (*β*2m), and peptide (pMHC-I) (3). The MHC-I contains α1, α2 and α3 domains in its extracellular region, with the α1 and α2 domains forming the pMHC-I peptide-binding groove (PBG) (4–9). Antigen peptides 8-11 amino acids in length are loaded onto the PBG and subsequently recognized by TCR expressed on the surface of CD8^+^ T cells (10). Interactions between the pMHC-I complex and TCR form the first signal for the activation of antigen-specific CD8^+^ T cells (11). The direct interaction between TCR and pMHC-I demonstrates the MHC-I-restriction nature of CD8^+^ T cell recognition and indicates a close evolutionary relationship between pMHC-I and TCR (12, 13). It is widely accepted that in addition to direct interaction with TCR, MHC-I engagement of the CD8 coreceptor is required for the functional activation of CD8^+^ T cells (14). The transmembrane glycoprotein CD8 molecules bind to pMHC-I and recruit the Src tyrosine kinase p65lck (Lck) to the TCR-pMHC-I complex, resulting in the assembly of a TCR-pMHC-I-CD8 ternary complex (15). Therefore, the presentation of various endogenous peptides by the pMHC-I complex is key in determining whether the antiviral CTL immune response is initiated.

The structures and functions of pMHC-I complexes that bind viral peptides or peptides have been investigated in humans (16), mice (17), monkeys (9), cattle (8), pigs (5), horses (18), dogs (6), cats (4), chickens (19), and a bony fish (20). The polymorphism of MHC-I molecules causes variation in PBGs in various mammalian pMHC-I complexes, which triggers various types of CTL immune responses. Although there are many reports of mammalian pMHC-I complexes, no study has yet been conducted on the pMHC-I structure in the bear family (Ursidae), which includes the giant panda. The bear family includes 5 genera and 8 species. Classical MHC-I genes and polymorphisms in the giant panda (*Ailuropoda melanoleuca*), brown bears and the Asiatic black bear have been reported. Phylogenetic studies indicate that although the giant panda belongs to the bear family, divergence between the giant panda and other bears occurred approximately >17 million years ago (21). The giant panda has an estimated population size of approximately 2500 individuals worldwide, and its classical MHC-I/II genes are located on chromosome 9q (22). The large genetic region of MHC-I is believed to play critical roles in CTL immunity. In addition, the giant panda MHC-I molecules represent an important model for understanding the structural and functional characteristics of these antiviral complexes in the bear family.

Several studies have been performed to identify giant panda MHC-I genes. In early work, three classical MHC-I genes (including *Aime-128*) were identified in the giant panda, and the *Aime-128* gene was proven to include ten conserved amino acids critical for viral antigenic peptide binding as described by human leukocyte antigen I (HLA-I) (23). Subsequent genetic analysis identified six panda MHC-I genes, four of which are classical MHC-I genes (24). High levels of genetic variation have been demonstrated in the panda MHC-I molecules (25), which is consistent with the selective advantage of MHC polymorphisms when encountering numerous types of pathogens (2). Pandas appear to be highly susceptible to zoonoses. Cases of canine distemper virus (CDV) and canine coronavirus (CCV) infections have occurred in panda populations (26, 27). Therefore, it is important to elucidate key molecules, such as the pMHC-I complex, and the antiviral CTL immune response of the panda, to aid vaccine development and the development of effective treatments for various diseases. In this report, the panda was chosen as a representative species of the bear family, and its pMHC-I structure (aka pAime-128) was determined by X-ray crystal diffraction. The PBG of panda pMHC-I has distinct features, from the amino acid sequence to the three-dimensional (3D) structure, and pAime-128 also reflects the features of the bear family. The findings reveal specific characteristics of the giant panda pMHC-I structure, greatly enhancing the understanding of MHC-I-related antiviral immunity in the bear family.

## RESULTS

### Structural Framework of Panda pAime-128

To verify the binding of these viral peptides to the giant panda pMHC-I (pAime-128), *in vitro* refolding experiments were carried out. The results revealed that four peptides can form stable pAime-128 complexes (**Supplementary Fig 1 and Table 1**). The complex of Aime-128 and Aime-*β*2m with the CCV-NGY9 peptide was crystallized in the P4_3_2_1_2 orthorhombic space group with a resolution of 2.68 Å (**Table 2**). The viral CCV-NGY9 peptide (CCV-NGY9) was selected from the CCV spike protein (S protein), which is among the structural proteins encoded by ORF2 (**Figure 1**). Similar to the S proteins of other coronaviruses, the S protein of CCV plays a fundamental role in the interaction with the cellular receptor and induces neutralizing antibodies in the natural host (28). Two complexes of pAime-128, molecule 1 (aka M1) and molecule 2 (aka M2), were found in one asymmetric unit. The root-mean-square deviation (RMSD) of the two molecules of Aime-128 was 0.584 Å. Because of the high consistency between M1 and M2, the following analysis of pAime-128 was based mainly on M1 (i.e., pAime-128 is M1 unless otherwise specified). The Aime-128 H chain contains the α1 (residues 1 to 90), α2 (residues 91 to 182) and α3 (residues 183 to 275) domains, of which the α1 and α2 domains constitute the PBG (**Fig 1A and B**). The α1 and α2 domains of the Aime-128 can be divided into two portions: one portion (α1, residues 57 to 84; α2, residues 138 to 150 and 152 to 176) forms helices located at the top of PBG, and the remaining portion (residues 3 to 13, 20 to 28, 31 to 37, 46 to 47, 93 to 103, 110 to 118, 121 to 126, and 133 to 135) forms an eight-stranded *β*-sheet platform at the bottom, named A to H. The CCV-NGY9 is loaded onto PBG of pAime-128 (**Fig 1B**).

**TABLE 1.** Peptides refolded with the Aime-128 and the Aime-*β*_2_m.

| Name | Derived protein | Start position | Sequence | %Random<sup>b</sup> | Refolding<sup>c</sup> |
| --- | --- | --- | --- | --- | --- |
| CCV-NGY9<sup>a</sup> | CCV spike | 436 | NGYNFFSTF | 0.064 | ++ |
| CDV-TSV9 | CDV H protein | 192 | TSVGRFFPL | 0.051 | ++ |
| CPV-NTY9 | CPV VP1 | 448 | NTYGPLTAL | 0.055 | ++ |
| H7N9-EVW9 | H7N9 | 428 | EVWSYNAEL | 0.166 | ++ |

<sup>a</sup> Peptide CCV-NGY9 was used in the structural determination of the giant panda Aime-128;
<sup>b</sup> % Random is a base value for estimating the binding affinities of peptides with the NetMHCpan 4.0 server (http://www.cbs.dtu.dk/services/NetMHCpan/): Rank threshold for strongly binding peptides, 0.500; rank threshold for weakly binding peptides, 2.000
<sup>c</sup> ++, peptide binds Aime-128 and can tolerate anion-exchange chromatography.

**TABLE 2.** X-ray diffraction data processing and refinement statistics.

| Statistic | Value for Aime-128/CCV-NGY9 |
| --- | --- |
| Data processing |  |
| Space group | $P4_32_12$ |
| Wavelength (Å) | 0.97892 |
| Cell dimensions (°) |  |
| $a, b, c$ (Å) | 173.68, 173.68, 84.35 |
|  | 90.0, 90.0, 90.0 |
| Resolution range (Å)<sup>a</sup> | 50.00-2.68 (2.75-2.68) |
| Total number of reflections | 193316 |
| Number of unique reflections | 34668 |
| Number of molecule in the asymmetric unit | 2 |
| Average redundancy<sup>a</sup> | 6.8 (7.2) |
| Completeness (%)<sup>a</sup> | 99.1 (100.0) |
| $R_{\text{merge}}$ (%)<sup>b</sup> | 9.4 (39.3) |
|  | 18.2 (10.3) |
| Refinement |  |
| Resolution (Å) | 36.62-2.68 |
| R-factor (%) | 19.0 |
| $R_{\text{free}}$ (%)<sup>c</sup> | 22.1 |
| RMS deviations from restraint target values: |  |
| Bond length (Å) | 0.004 |
| Bond angles (°) | 0.841 |
| Average B factor | 27.7 |

<sup>a</sup> Numbers in parentheses indicate the highest-resolution shell.
<sup>b</sup> $R_{\text{merge}} = \sum h \sum I_{ih} - \langle I_h \rangle / \sum h \sum I \langle I_h \rangle$ , where $\langle I_h \rangle$ is the mean intensity of the observation $I_{ih}$ of reflection $h$ .
<sup>c</sup> $R$ factor = $\sum (F_{\text{obs}} - F_{\text{calc}}) / \sum F_{\text{obs}}$ ; $R_{\text{free}}$ is the $R$ factor for a subset (5%) of reflections that was selected prior to refinement calculations and not included in the refinement.

**FIG 1.**
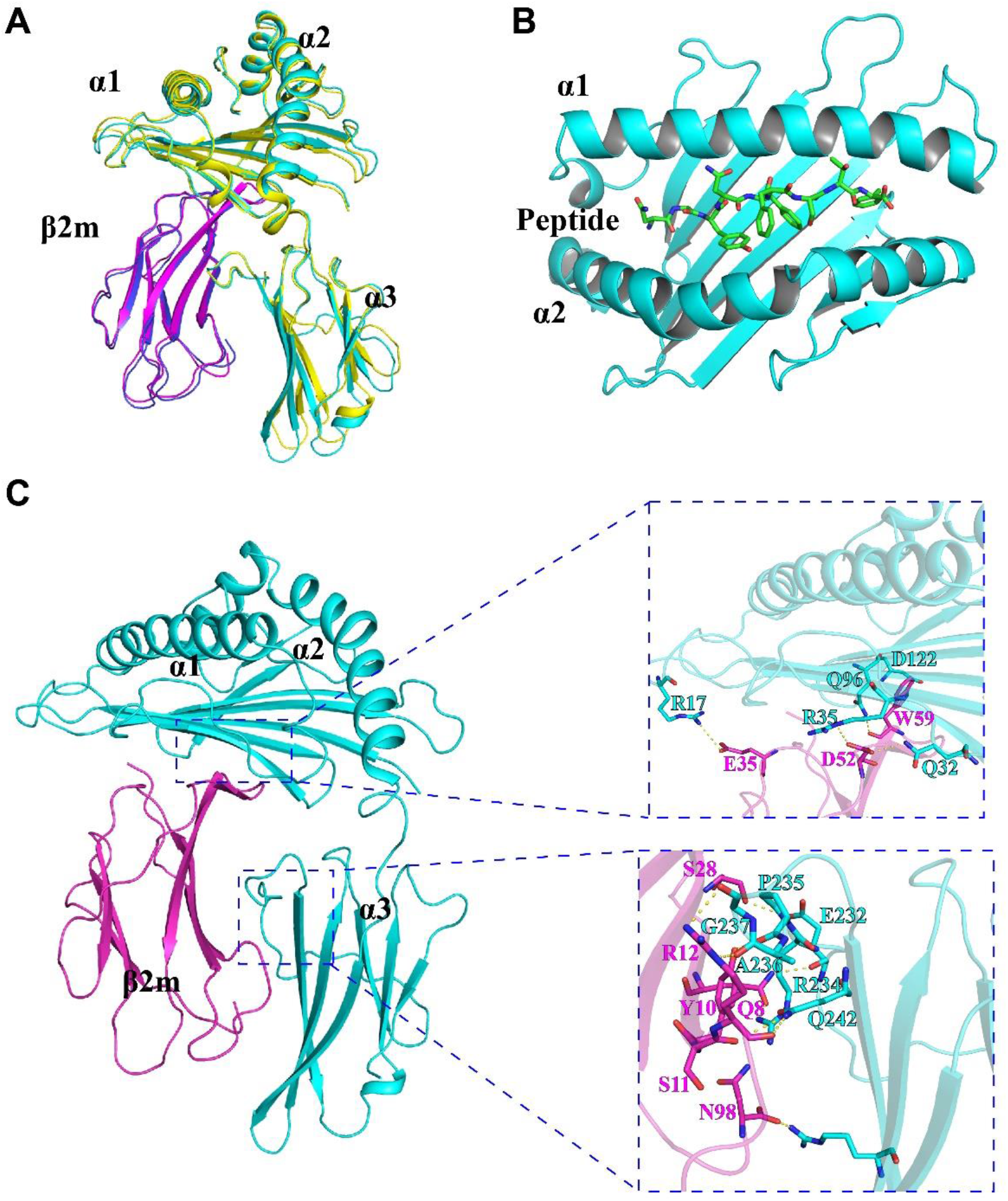
Structure of pAime-128 complex. (**A**) Overview of M1 and M2 in an asymmetric unit of pAime-128 complex. Aime-128 M1 and M2 are indicated in cyan and yellow, respectively. (**B**) The formation of the CCV-NGY9 peptide in PBG of pAime-128. (**C**) Interactions between Aime-128 and Aime-*β*_2_m. Aime-128 is shown in cyan and Aime-*β*_2_m is shown in magenta. The hydrogen bonds formed between Aime-128 and Aime-*β*_2_m are shown as yellow dashed lines.

Further analysis showed that strands and loops of Aime-*β*2m broadly interact with Aime-128, and the numbers and patterns of hydrogen bonds formed in these interactions were similar to those of other mammalian MHC I molecules. Analysis revealed that a depth of Aime-β2m in Aime-128 up to 1351.1 Å^2^ (**Fig 1C**). In addition, the hydrogen bonds involved in the interactions between Aime-128 and Aime-*β*2m were found to differ from those in humans. Aime-*β*2m forms only 15 hydrogen bonds with Aime-128, whereas in HLA-B*5101, *β*2m can form 25 hydrogen bonds with MHC-I (29), which implies the interactions are weaker than humans (**Fig 1C**). Thus, the main function of *β*2m is to stabilize MHC-I, even though the same *β*2m can interact with MHC-I in various ways. The main role of β2m is to ensure the stable function of pMHC-I, and the manner of its interaction with MHC-I is thought to be able to promote side effects such as *β*2m dissociation and even diseases (29).

### The Unique Pockets for Antigen-binding Peptide Found in pAime-128

The amino acids in PBG form six functional pockets, named pockets A to F, to accommodate residues of the CCV-NGY9 peptide (**Fig 2**). Pocket A in pAime-128 consists of residues Met^5^, Tyr^7^, Tyr^59^, Arg^62^, Asn^63^, Tyr^159^, Glu^163^, Trp^167^ and Tyr^171^ (**Table 3**). P1-Asn of CCV-NGY9 was inserted into pocket A. Based on the 3D structures of pMHC-I in different species, Asn^63^ and Glu^163^ of Aime-128 are rare among all of the known class I molecules, i.e., these residues are found in only certain alleles, such as HLA-B*5101 and HLA-B*2705 (**Fig 2**). Due to the presence of residue Asn^63^ in Aime-128, pocket A of this molecule is considered to be tighter than that of the other pMHC-I complexes (**Fig 3**).

**TABLE 3.** Hydrogen bonds and van der Waals interactions between peptides and Aime-128

| Hydrogen Bonds and Salt Bridge |  |  |  |  |
| --- | --- | --- | --- | --- |
| Peptide CCV-NGY9 |  | Aime-128 |  |  |
| Residue | Atom | Residue | Atom | Van der Waals Forces |
| P1-Asn | N | Tyr<sup>171</sup> | OH | Met<sup>5</sup>, Tyr<sup>7</sup>, Tyr<sup>59</sup>, Arg<sup>62</sup>, Asn<sup>63</sup>, Ile<sup>66</sup>, Trp<sup>167</sup>, Tyr<sup>171</sup><br>(63) |
|  | O | Tyr<sup>159</sup> | OH |  |
| P2-Gly | N | Asn<sup>63</sup> | OD1 | Tyr<sup>7</sup>, Asn<sup>63</sup>, Ile<sup>66</sup>, Tyr<sup>159</sup> (23) |
|  |  | Tyr<sup>7</sup> | OH |  |
| P3-Tyr | OH | Glu<sup>152</sup> | OE1 | Tyr<sup>9</sup>, Ile<sup>66</sup>, Asn<sup>70</sup>, Trp<sup>97</sup>, His<sup>99</sup>, Glu<sup>152</sup>, Arg<sup>155</sup>,<br>Tyr<sup>156</sup>, Tyr<sup>159</sup> (91) |
|  |  | Arg<sup>155</sup> | NE |  |
| P4-Asn | O | Arg<sup>155</sup> | NH1 | Ile<sup>66</sup>, Asp<sup>69</sup>, Asn<sup>70</sup>, Arg<sup>155</sup> (27) |
|  | O | Arg<sup>155</sup> | NH2 |  |
| P5-Phe |  |  |  | Asn<sup>70</sup>, Ala<sup>73</sup>, Trp<sup>97</sup>, Trp<sup>147</sup>, Glu<sup>152</sup>, Arg<sup>155</sup> (36) |
| P6-Phe |  |  |  | Glu<sup>152</sup>, Arg<sup>155</sup> (16) |
| P7-Ser | OG | Glu<sup>152</sup> | OE2 | Trp<sup>147</sup>, Ala<sup>150</sup>, Glu<sup>152</sup> (19) |
| P8-Thr | O | Trp<sup>147</sup> | NE1 | Ala<sup>73</sup>, Val<sup>76</sup>, Asp<sup>77</sup>, Thr<sup>143</sup>, Trp<sup>147</sup> (30) |
| P9-Phe | N | Asp<sup>77</sup> | OD1 | Asp<sup>77</sup>, Thr<sup>80</sup>, Tyr<sup>84</sup>, Ile<sup>95</sup>, Leu<sup>116</sup>, Tyr<sup>123</sup>, Thr<sup>143</sup>,<br>Lys<sup>146</sup>, Trp<sup>147</sup> (84) |
|  | O | Tyr<sup>84</sup> | OH |  |
|  | O | Thr<sup>143</sup> | OG1 |  |
|  | O | Lys<sup>146</sup> | NZ |  |

<sup>a</sup> Numbers in parentheses are the amounts of van der Waals force.

**FIG 2.**
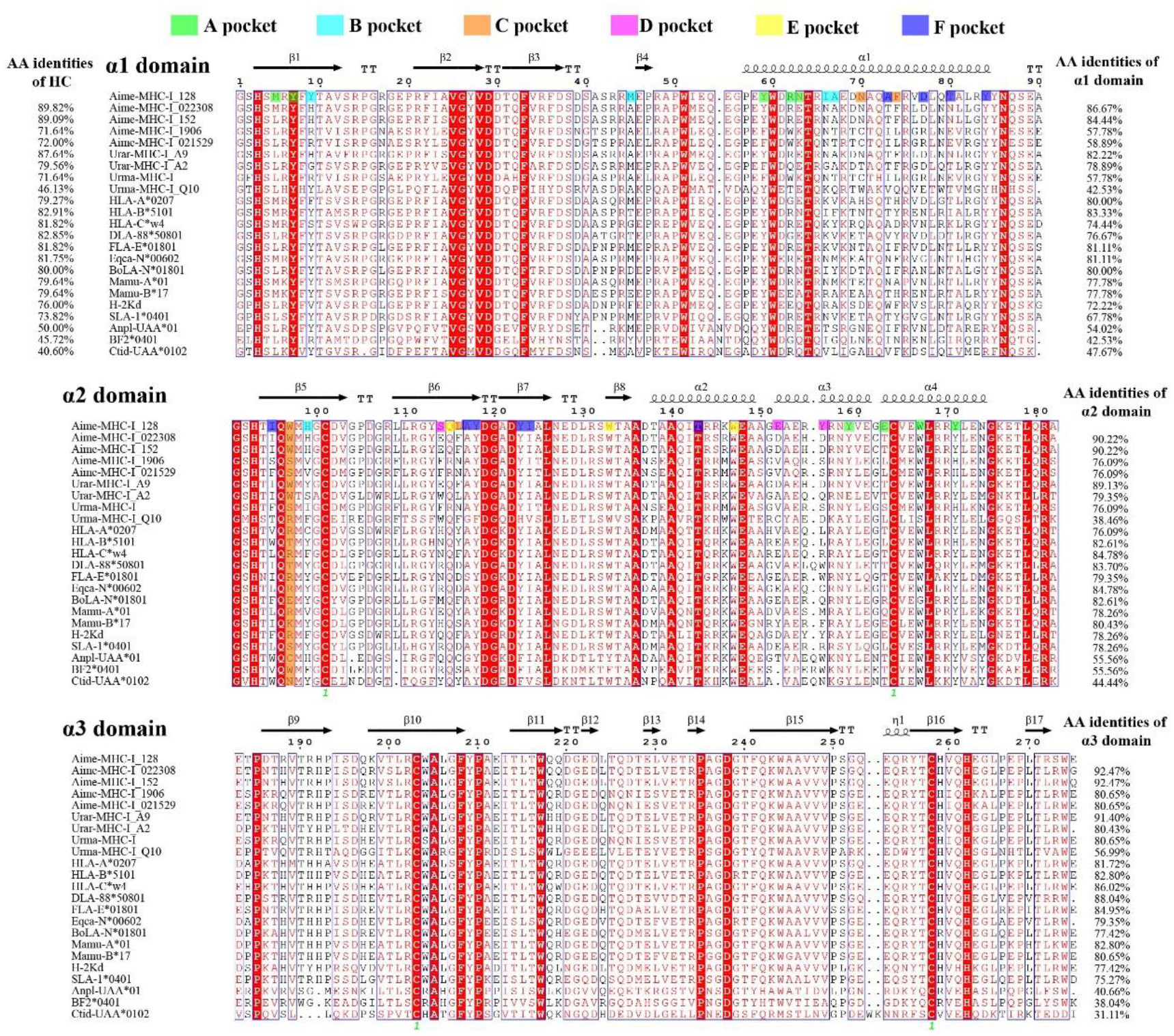
Structure-based amino acid sequence alignment of Aime-128 and other representative mammalian MHC-I molecules with available crystal structures. Cylinders indicate α helices, and black arrows above the alignment indicate *β* strands. The amino acid identities of different domains between Aime-128 and the listed MHC-I molecules are given on the right-hand side. The residue positions contributing to Aime-128 pockets are highlighted by pocket-specific colored shading.

**FIG 3.**
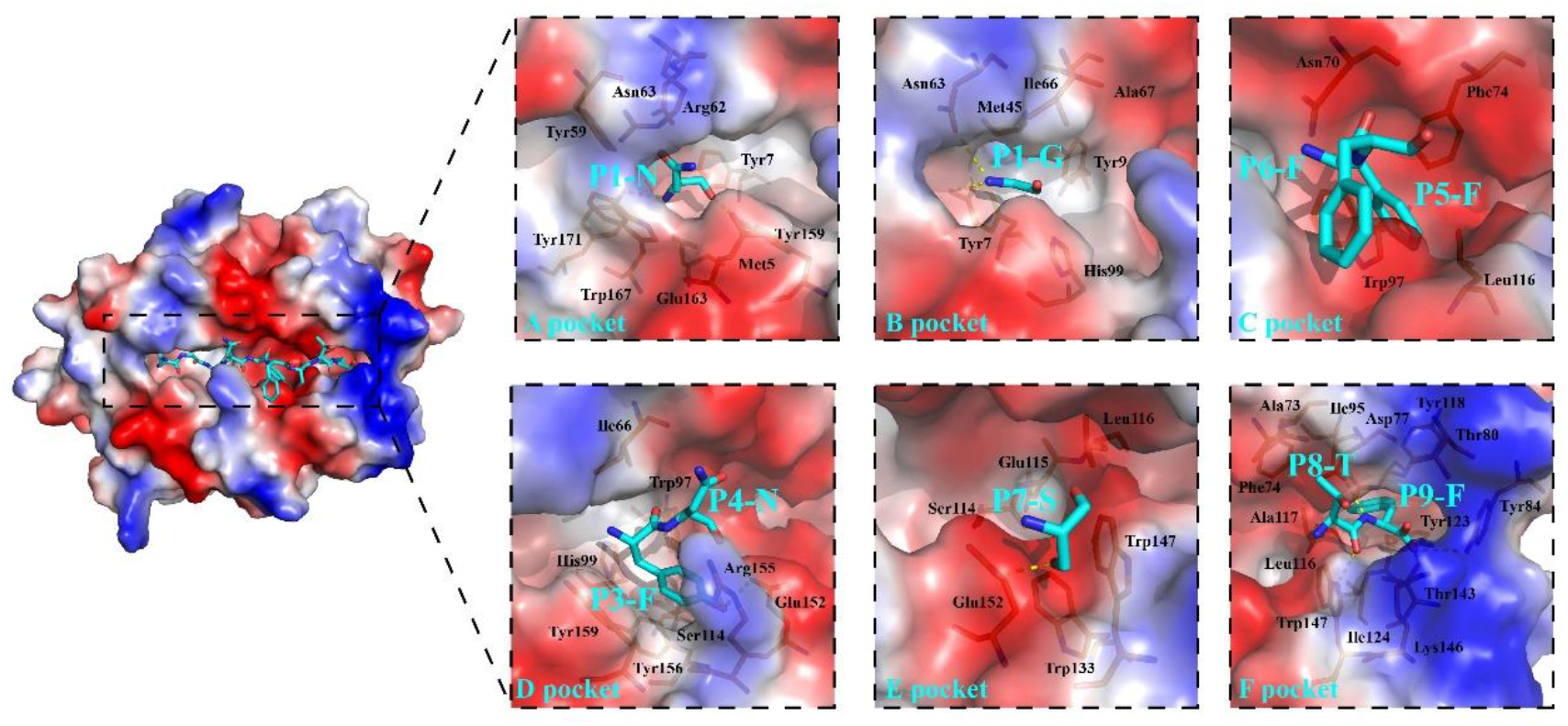
Structural analysis and comparison of the six pockets of pAime-128 complex. The six pockets are shown in surface charge representation (blue, positively charged; red, negatively charged; white, nonpolar). The pocket residues were determined based on interaction with peptide ligand as indicated by CCP4 software and on our visual inspection of pocket continuity. The residues in PBG of pAime-128 complex are shown in stick form. The residues of CCV-NGY9 are shown as cyan sticks, and hydrogen bonds between peptide and pockets are shown as yellow dashed lines.

Residues Tyr^7^, Tyr^9^, Ala^24^, Met^45^, Asn^63^, Ile^66^, Ala^67^ and His^99^ form pocket B of PBG in pAime-128 (**Fig 3 and Table 3**). P2-Gly of CCV-NGY9 is tethered by hydrogen bonds to residues in PBG and fits into the small pocket B. In most class I molecules, the residues at positions 63 and 66 are Glu and Lys, respectively, which are both charged residues. The residue at position 99 of Aime-128 is His instead of the conserved Tyr residue; both of these residues have a large side chain, but they have different charges (**Fig 2**). Pocket B of pAime-128 extends deep under the α1 helix and touches the Met45 residue, which is one of the most polymorphic residues in mammalian MHC-I molecules (**Fig 3**).

Pocket C of pAime-128 is composed of residues Asn^70^, Phe^74^, Trp^97^ and Leu^116^ and shows an obvious negative charge, and none of these residues are highly conserved among known mammalian class I molecules (**Fig 2**). Pocket C is wide but shallow. P5-Phe and P6-Phe of CCV-NGY9 are located in this pocket, and no hydrogen bonds exist between the C pocket and the peptide, only van der Waals forces. The side chain of P6-Phe extends outward into the solvent and may play a role in TCR docking (**Fig 3**).

Pocket D in pAime-128 accommodates the side chains of P3 and P4 residues. The pocket consists of residues Ile^66^, Trp^97^, His^99^, Ser^114^, Glu^152^, Arg^155^, Tyr^156^ and Tyr^159^ (**Table 3**). In the peptide CCV-NGY9, the residue at position 3 is Tyr, which has a large side chain and, as a result, forms two hydrogen bonds and many van der Waals interactions with pocket D (**Fig 3**). The side chain of P3 is inserted into pocket D and forms hydrogen bonds with Glu^152^ and Arg^155^. Notably, Glu^152^ and Arg^155^ are not conserved in most mammalian class I molecules, and both of these residues are charged (**Fig 2**). P3-Tyr is primarily surrounded by the residues Trp^97^, His^99^, Tyr^156^ and Tyr^159^, which have large side chains and provide a clear boundary to pocket D. The small residue Ser^114^ provides enough space in the pocket to accommodate the large side chain of P3-Tyr (**Fig 3**). The side chain of P4-Asn extends outward into the solvent for potential recognition by TCRs. Different formations of hydrogen bonds between the PBG and the peptide were also observed for peptide residues P3-Tyr and P4-Asn.

Pocket E is composed of residues Ser^114^, Gln^115^, Leu^116^, Trp^133^, Trp^147^ and Glu^152^ (**Fig 2 and Table 3**). The pocket is obviously negatively charged and located relatively deep beneath the α2 helix, and residues Ser^114^, Gln^115^ and Leu^116^ constitute the bottom of platform. P7-Ser of CCV-NGY9 is inserted into pocket E (**Fig 3**).

Pocket F of pAime-128 PBG consists of the highly conserved residues Ala^117^, Tyr^118^, Tyr^123^, Ile^124^, Thr^143^, Lys^146^ and Trp^147^, as well as the poorly conserved residues Ala^73^, Phe^74^, Asp^77^, Thr^80^, Ile^95^ and Leu^116^ (**Fig 2**). Among these residues, Leu116 is not present in any other annotated MHC class I molecule deposited in the Protein Data Bank (18, 19). Pocket F is therefore narrow but deep enough to accommodate large residues, and residues P8-Thr and P9-Phe are located in this pocket (**Fig 3**). The anchor residue P9-Phe is similar to the anchor residues of HLA-A*01, HLA-B*35, and HLA-B*57, i.e., large residues with an aromatic ring (30). The aromatic ring in CCV-NGY9 is held in close contact with residues in pocket F by strong hydrogen bonds and van der Waals contacts (**Fig 3**).

### Comparison of panda Aime-128 to Other Known MHC-I Molecules

The amino acid homology of MHC-I molecules in giant pandas is approximately 70-90%, but the homology of the peptide-binding domains composed of the α1 and α2 regions is low (**Fig 3**). These results suggest that the antigen-peptide presenting profiles of the panda individuals are different, especially the homology of the α1 region, which can be as low as approximately 57%; thus, the difference is large. The results also show the potential differences in pMHC-I complexes among the bear family. It can be concluded that classical MHC-I binding antigen-peptide profiles of the bear family are diverse.

Structure-based amino acid sequence analysis implied that MHC-I molecules may form different PBGs in pandas. However, the amino acid composition of the A, B, C, D, E and F pockets in the PBG was not identical among pandas, and the PBG regions of the pandas differed extensively from those of other members of the bear family (**Fig 3**). Of the ten conserved amino acids in the α1 and α2 domains of Aime-128 predicted to be critical for peptide binding (23), only eight (Met^5^, Tyr^7^, Tyr^59^, Tyr^84^, Thr^143^, Lys^146^, Tyr^159^ and Tyr^171^) were confirmed in our study to have direct contact with the CCV-NGY9 peptide (**Table 3**). In addition, the amino acid composition of PBGs among other genera and members of the bear family is not completely conserved, so there are some differences in pockets A to F. There are also great differences among giant pandas; other members of the bear family; other mammals; such as humans and mice; and nonmammals, such as the bony fish grass carp (**Fig 3**).

### Validation of Anchor Sites and Binding Motif in pAime-128

The peptide binding to pAime-128 PBG used in a featured mode with a central bulge, as revealed by unambiguous electron density (**Fig 4A**). Extensive comparison of pAime-128 with other pMHC-I structures deposited in the Protein Data Bank showed that the main chain of the CCV-NGY9 peptide has a characteristic conformation similar to that presented by HLA-B*0801 (**Fig 4B**). Ternary TCR-peptide-HLA-B*0801 structure of epitope HCV-HSK9 (HSKKKCDEL, hepatitis C virus-derived) showed that the protruding residue Lys-4 in HCV-HSK9 plays important roles in TCR recognition (31, 32). In pAime-128, the side chain of P6-Phe pointed upward for solvent exposure, similar to the side chain of Lys-4 in HCV-HSK9 (**Fig 4A and B**).

**FIG 4.**
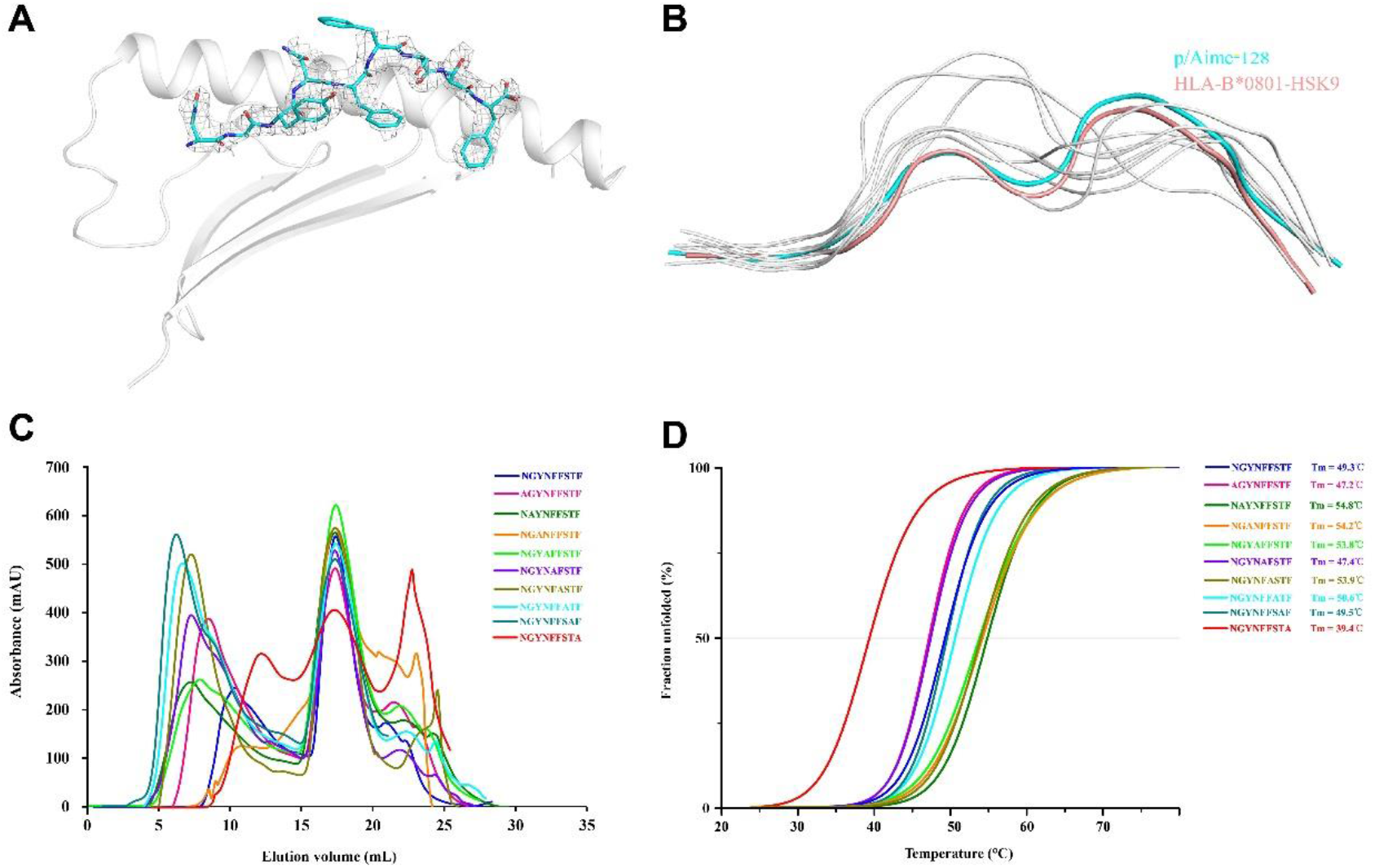
CCV-NGY peptide conformations for TCR contacts. (**A**) Electron densities and overall conformations of the structurally defined CCV-NGY9 peptides of pAime-128. (**B**) Superposition of the CCV-NGY9 peptide presented by pAime-128 with nonapeptides presented by other vertebrate MHC-I molecules. Peptides are shown as ribbons with the following color scheme: cyan, Aime-128-CCV-NGY9; salmon, HLA-B*0801-HSK9 (4QRP); light blue, HLA-B*0801-FLR9 (1MI5); and gray, peptides presented by HLA-A*0201 (2AV1), Mamu-A*02 (3JTT), H-2Kd (1VGK), SLA-1*0401 (3QQ3), BoLA-A*01801 (3PWU), FLA-E*01801 (5XMF), DLA-88*50801 (5F1I) and *Anpl*-UAA*01 (5GJX). (**C**) The refolded products of Aime-128 and Aime-*β*2m in the presence of mutated peptides tested by gel filtration chromatograms. The refolding efficiencies are represented by the relevant concentration ratios and by the heights of the peak for each mutant. The mutated peptides P1A, P5A and P9A clearly yielded low refolding efficiency. (**D**) CD spectropolarimetry was utilized to assess the thermostabilities of the purified pAime-128 complexes. Shown here are the data fitted to the denaturation curves using the Origin 9.1 program (OriginLab). The Tm values of different peptides are indicated by the gray line at the 50% unfolded level.

With different peptide presentation modes observed for pAime-128, it is necessary to clarify the critical anchors for the peptide bound to Aime-128. Therefore, the stabilities of the CCV-NGY9 peptide and its mutants with alanine substitution at each position bound with Aime-128 were measured through circular dichroism (CD) spectroscopy (**Fig 4C and D**). The midpoint transition temperature (Tm) value of the wild-type CCV-NGY9 peptide was determined to be 49.3°C. Compared with the CCV-NGY9 peptide, the CCV-P9A-substituted peptide showed a dramatic decrease in stability with the lowest Tm at 39.4°C, while the peptide mutants CCV-P1A and CCV-P5A had lower Tms of 47.2°C and 47.4°C, respectively, indicating a minor decrease in stability. The remaining peptide mutants showed different levels of enhanced stability, with Tms ranging from 49.5°C to 54.8°C (**Fig 4C and D**). These data indicated that for peptides bound to Aime-128, PΩ plays a pivotal role in peptide anchoring since it cannot be replaced by alanine. The peptide residues at the N terminus (PN) and position 5 (P5) are also important for stabilization of pAime-128 complex. Notably, the mutant CCV-P2A displayed the highest Tm of 54.8°C, suggesting that the mutation of P2-G to comparatively large amino acids might increase the stability of pAime-128 complexes. Based on this finding and the peptide screening results (**Table 1**), we identified the motif of pAime-128 presented peptides. The binding motif of pAime-128 is (Ala, Asn, Thr or Glu)-(Ala, Gly, Ser, Thr or Val)-x-x-(Ala, Phe, Arg, Pro, Tyr)-x-x-x-(Phe or Leu).

### Distinctive Interaction of Aime-128 and *β*2m/TCR/CD8 Molecules

Superposing pAime-128 molecule on the known pMHC-I structures from different species showed that the AB loop of pAime-128 and other mammal pMHC-I structures does not contact *β*2m (**Fig 5A-E**), whereas the AB loop binds *β*2m via hydrogen bonds only in nonmammals (**Fig 5F, G**). Furthermore, the CD and EF loops of mammals are longer than those of nonmammals (**Fig 5A**), because mammalian MHC-I contains two more amino acids than nonmammalian MHC-I (**Fig 2**).

**FIG 5.**
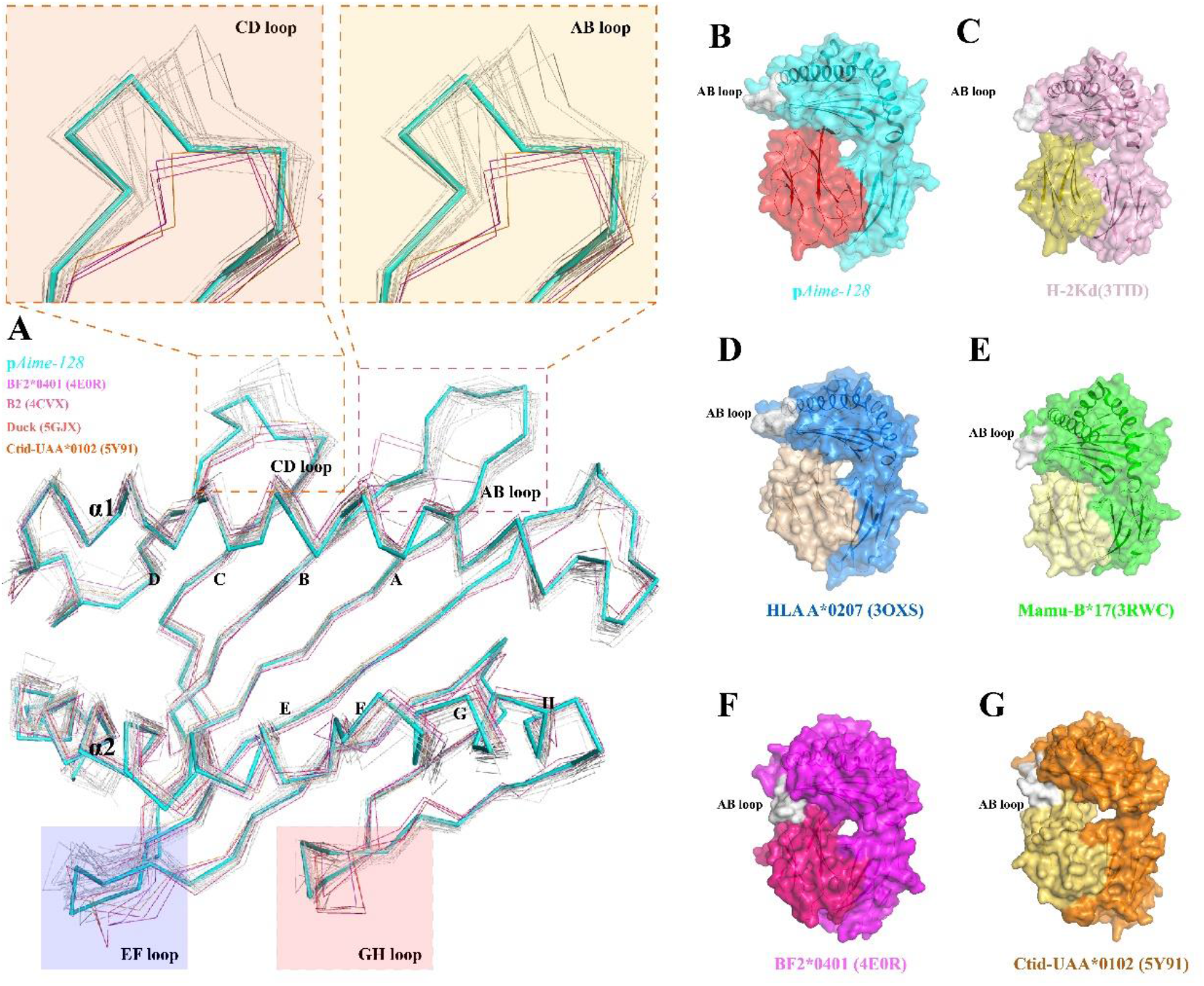
pAime-128 complexes show some features that are similar among mammals but different from those of lower vertebrates. (**A**) Unique details of the higher vertebrates in PBG. (pAime-128 cyan, grass carp-5Y91 light orange, chicken-4E0R hot pink, duck-5GJX magenta and other mammals (1VGK, 2XFX, 3OXS, 3PWU, 3TID, 1ZVS, 3QQ3, 3RWC, 3X11, 4QRQ, 4MJ6, 4MNX, 4N02, 4NT6, 4O2C, 4QOK, 4QRS, 5XMF, 3BUY, 1ZT7, 4WJ5 white). The AB loops of the nonmammals are downward, which can interact with *β*2m, while the mammals’ cannot contact *β*_2_m. The CD loops of mammals are longer than those of lower vertebrates, causing the 2 more residues after the 40^th^ amino acids of PBG in the higher vertebrates than in the nonmammals. The EF loops of mammals are longer than those of nonmammals. There is a hydrophobic core in the GH loop of grass carp, duck and chicken, so the GH loop is expanded and close to *β*_2_m. (**B-G**) The gray areas indicate the discrepant regions of MHC-I and *β*_2_m interfaces among chicken, grass carp, and mammals. (**B**) In pAime-128, the AB loop of Aime-128 cannot bind to *β*_2_m. (**C**) In mouse pMHC-I structures (3TID), the AB loop cannot bind to *β*_2_m. (**D**) In human pMHC-I structures (3OXS), the AB loop cannot bind to *β*_2_m. (**E**) In monkey pMHC-I structures (3RWC), the AB loop cannot bind to *β*_2_m. (**F**) In chicken, the BF2*0401(4E0R) AB loop can bind to *β*_2_m. (**G**) In grass carp (5Y91), the AB loop can bind to *β*_2_m.

The antigenic peptides presented by pMHC-I complexes are eventually scanned and recognized by TCRs to initiate MHC-I-restricted CD8 T cell immunity. Extensive structural studies have illustrated that the TCR repertoire specific for a certain antigenic peptide is determined by the conformations of both the peptide and the restricted MHC-I element (12, 33). Comparisons with MHC-I structures with known TCR docking strategies revealed that the exposed Aime-128 residues Arg^62^, Arg^65^, Ile^66^, Asp^69^, Gln^72^, Val^76^, Gln^79^, Glu^154^, Arg^155^, Asn^158^, Glu^161^, Glu^163^, Glu^166^ and Trp^167^ have great potential to interact with TCRs (**Fig 6A**). Although most of these residues are highly conserved or preferentially used among different species, rarely used residues were observed in Aime-128, including Ile^66^, Asp^69^, Arg^155^ and Asn^158^ (**Fig 2**).

**FIG 6.**
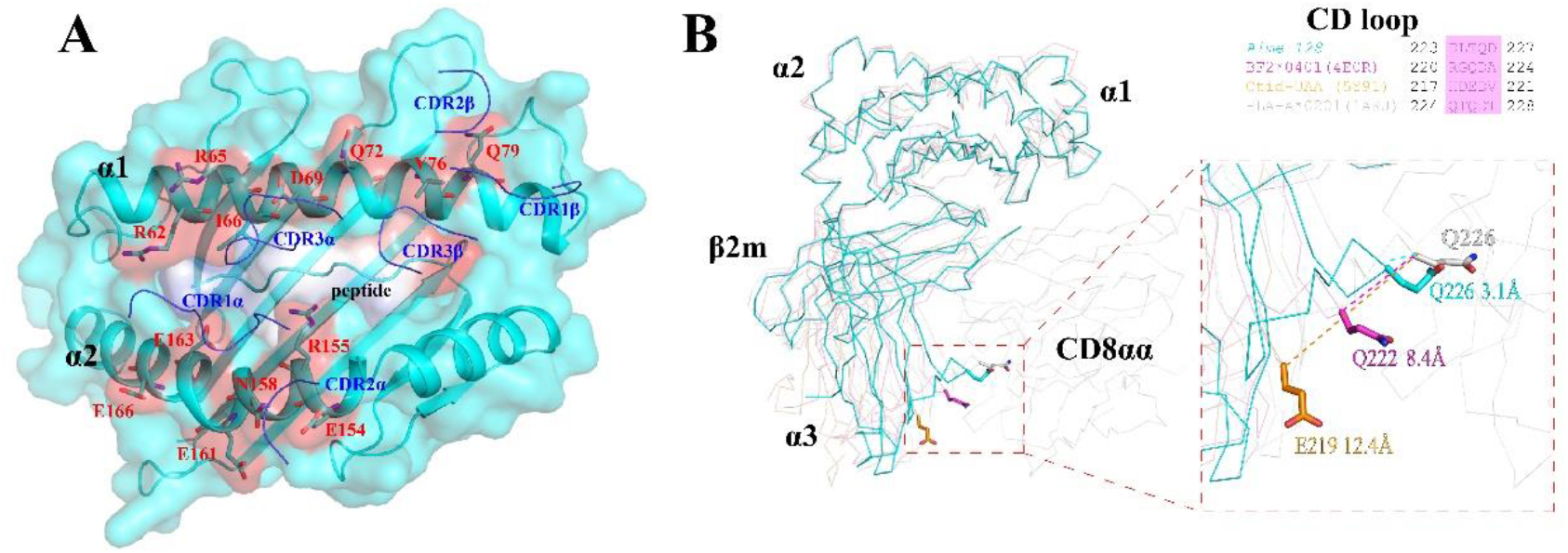
Putative TCR docking sites on pAime-128 complexes and a unique way to bind CD8. (**A**) Based on the HLA-B*0801-HSK9-specific TCR, the residues on pAime-128 that may contact the TCR CDR loops (green) are shown as red surfaces, and the CCV-NGY9 peptide is shown on the surface according to its charge. The proximity of the TCR CDR loops to the peptide-binding region of pAime-128 is shown. (**B**) The major shift in the α3 domain and the variation in the key residues for binding CD8αα in pAime-128 complexes. pAime-128 structure is superposed on the HLA-A2-CD8αα (HLA-A2, white, PDB code: 1AKJ), *Ctid*-UAAg (cyan, PDB code: 5Y91) and BF2*0401 (hot pink, PDB code: 4E0R) structures in the ribbon. The residues shown in different colors according to their species in the CD loop that are critical for interaction with CD8αα are shown in sticks. The distance between E219 of p*Ctid*-UAAg and Q226 of HLA-A*0201 (PDB: 1AKJ) is approximately 12.4 Å, and the distance between Q222 in BF2*0401 (PDB: 4E0R) and Q226 of HLA-A*0201 (PDB: 1AKJ) is approximately 8.4 Å. The distance between the superposed CD loops of pAime-128 is approximately 3.1 Å.

Further analysis found different structures of the α3 loop (CD loop) in pAime-128 and other pMHC-I molecules. The CD loop of Aime-128, encompassing residues 222-228, is believed to be essential for CD8 interaction (34). Of particular note, the highly conserved residues Glu^222^ and Gln^226^, which have been proven to directly contact CD8 (35), are also present in pAime-128. Interestingly, the distance between Glu^219^ of p*Ctid*-UAAg and Gln^226^ of HLA-A*0201 is approximately 12.4 Å, and the distance between Gln^222^ in BF2*0401 and Gln^226^ of HLA-A*0201 is approximately 8.4 Å. The distance between the superposed CD loops of pAime-128 and HLA-A*0201 is approximately 3.1 Å (**Fig 6B**).

### Analysis of Peptide-epitopes of Viruses Based on the Motif of Panda pAime-128

The amino acid sequences of the pathogenic viruses related to viral diseases reported in the giant panda were obtained. Based on the 3D structure of pAime-128, the characteristics of presentation of viral peptides, and the motif from the experiment, the potential viral CTL epitopes of canine distemper virus (CDV), canine corona virus (CCV), giant panda polyomavirus (GPPV), giant panda rotavirus (GPRV), giant panda-associated gemycircularvirus (GPGE), giant panda anellovirus (GPAN), and influenza H1N1 viruses (36, 37) were reasonably proposed. The results are shown in **Fig. 7**. A series of viral epitope-peptides from the proteomes of viruses, including CCV, CDV, GPP, GPRV, GPGE, GPAN and H1N1, were screened against the panda MHC-I molecules. We obtained a total of 8 nonapeptides conforming to virus peptide motif of Aime-128, and synthesize them for *in vitro* verification. We found that all the selected virus peptides can bind to Aime-128, only one can bind Aime-128 but cannot tolerate anion-exchange chromatography while the others can tolerate anion-exchange chromatography. Although the result was rational calculated, it also fully demonstrates the potential research value of pAime-128.

**FIG 7.**
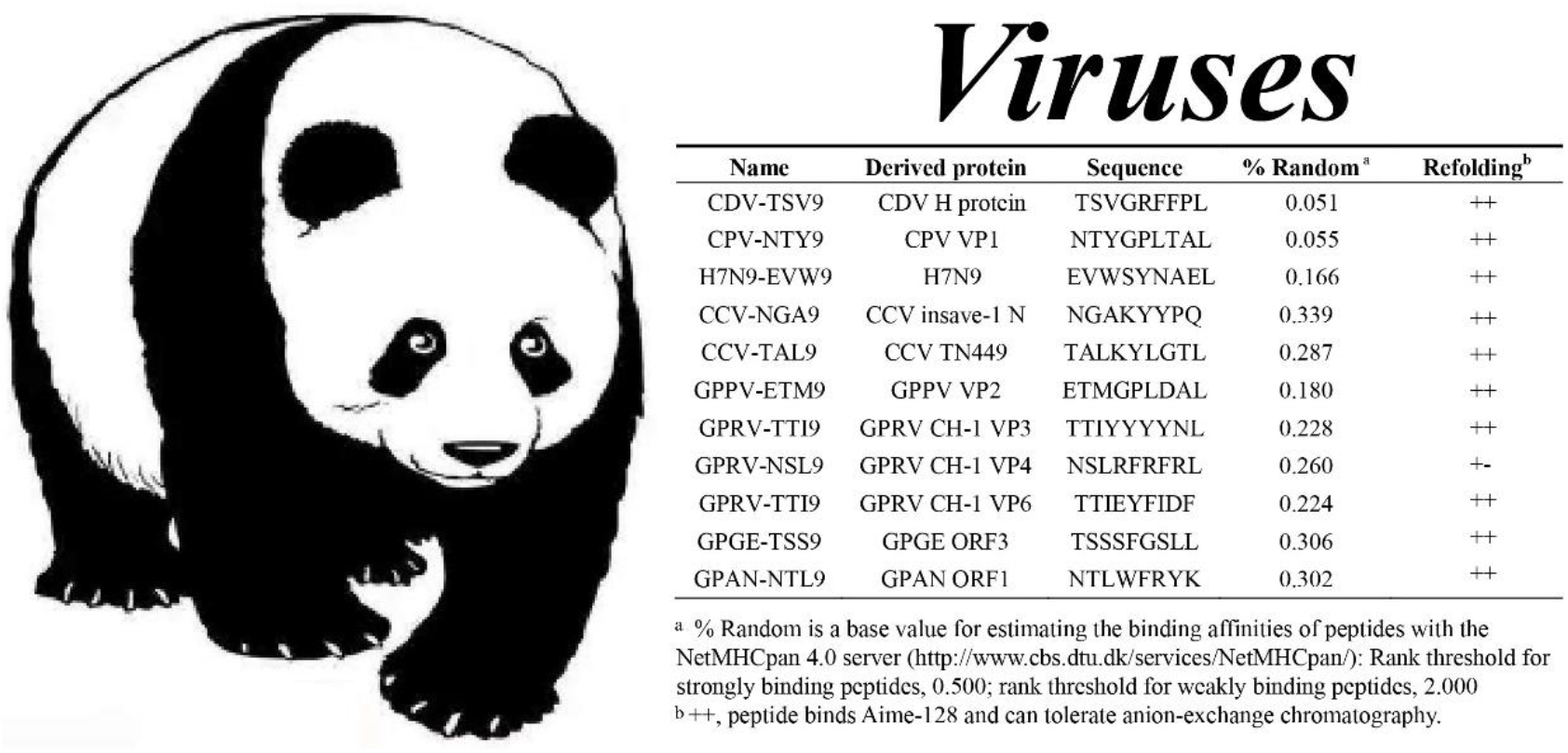
Predicted peptides and their binding to Aime-128 evaluated by *in vitro* refolding.

## DISCUSSION

The viral peptides presented by classical MHC-I molecules require the assembly of a pMHC-I complex for TCR recognition, which is critical for the initiation of antiviral CTL immunity in most jawed vertebrates (38). In this study, the 3D structure of giant panda classical MHC-I complexed with viral peptide derived from CCV spike protein was determined for the first time, and several unique features of panda pMHC-I were identified․.

One notable finding was that the homologies of classical MHC-I molecules in pandas are high, at approximately 70-90%", are higher than those of other mammals, being approximately 70-90%, whereas the homology of the antigen-binding domains composed of α1 and α2 regions is much lower. This result implied that the panda MHC-I molecules can form different PBGs. In addition, the amino acid composition of the A, B, C, D, E and F pockets in the PBGs are not identical among pandas, and the PBG regions of pandas and other members the bear family are very different from one another (**Fig 2**). The results suggest that the antigen-peptide presenting profiles of the panda individuals are different, especially the homology of the α1 region, which can be as low as approximately 57%. This difference may be quite remarkable. Thus, the classical MHC-I binding antigen-peptide profile of pandas is diverse. These findings indicate the high-throughput antiviral CTL immune response potential of Ursidae.

A second notable finding is that in the PBG pockets, the 9 residues constituting pocket A can be found in most giant panda MHC-I molecules, though 2 residues, Asn^63^ and Glu^163^, are rare among known MHC-I molecules. Due to the presence of the Asn^63^ residue, the pAime-128 complex pocket A is reasonably tight. The 8 residues that constitute pocket B can be found in almost all classical panda MHC-I molecules, although Asn^63^, Ile^66^, Ala^67^ and His^99^ are rare among known panda MHC-I molecules. Moreover, 5 residues constitute pocket C, but these residues are not highly conserved in mammalian MHC-I molecules (2). Similarly, 5 residues constitute pocket D, but Glu^152^ and Arg^155^ are not conserved. Therefore, although some amino acids are highly conserved in the composition of certain pockets, the different amino acids result in distinctive pocket conformations and a unique pAime-128 PBG for presenting viral epitope peptides with specific characteristics. Due to pAime-128, the polymorphism of amino acid sequences is easily reflected in its 3D structure, which is also important for studying pMHC-I complexes. In addition, the amino acid composition of PBGs among other genera and members of the bear family is not completely conservative, so there are some differences in A to F pockets. There are also great differences between giant pandas and other members of the bear family. The results also reveal the characteristics of the pMHC-I complexes of pandas and other members of the bear family and differences between them in their viral antigen-presenting peptide profiles.

Furthermore, by analyzing the structure of pAime-128 and comparing it with the structures of other known pMHC-Is, we found that the viral peptide in PBG features the “M” configuration that can activate T cells (45) and that key for recognition by TCR is the combination of the peptide P4-Asn and P6-Phe amino acids. Through analysis of the pockets in the PBG and the stability of CCV-NGY9 peptide and its alanine substitution mutant determined by CD spectroscopy, we identified the peptide-binding motif of pAime-128 and obtained a series of viral peptides that can bind or can potentially bind with pAime-128. The results provide a new platform for the effective design of bear family viral vaccines.

In summary, the structure of pAime-128, as a representative of the bear family pMHC-I complexes, was elucidated, and unique features of pMHC-I antigen presentation in in the bear family were identified. In addition, the viral peptide presentation profile was proposed in this paper. These results provide a new platform for the further study of bear family antiviral CTL immunology and vaccinology.

## MATERIALS AND METHODS

### Viral Peptide Synthesis

Four epitope peptides that potentially bind to Aime-128 were predicted by the NetMHCpan4.0 server (http://www.cbs.dtu.dk/services/NetMHC/) and via artificial intelligence selection based on the CCV spike protein (GenBank accession no. AFG19726.1), the CDV hemagglutinin (H) protein (GenBank accession no. AAD54600.1), CPV virus protein 1 (VP1) (GenBank accession no. AAV36771.1) and the H7N9 influenza A virus (GenBank accession no. AGI60301.1). These peptides were synthesized and purified by reverse-phase high-performance liquid chromatography (HPLC) (SciLight Biotechnology, Beijing, China) with >90% purity. The peptides were stored in lyophilized aliquots at −20°C after synthesis and were dissolved in dimethyl sulfoxide (DMSO) before use.

### Preparation of Expression Constructs

DNA fragments encoding the extracellular domain of giant panda MHC-I *Aime-128* (GenBank: AM282693, residues 1–270 of the mature protein with *Nde*I and *Xho*I restriction sites) and *Aime-β2m* (GenBank: AB178590, residues 1–98 of the mature protein with restriction sites *Nde*I and *Xho*I) were synthesized by Invitrogen Life Technologies (Shanghai, China). The products were ligated into the pET21a vector (Novagen) and then transformed into the *Escherichia coli* (*E. coli*) Transetta (DE3) strain. Recombinant plasmids were expressed as inclusion bodies and purified as previously described (29). Complexes of the CCV-NGY9 peptide with Aime-128 and Aime-*β*2m (pAime-128) were prepared with refolding assays using the gradual dilution method, as previously described (18). After 48 hours of incubation at 4°C, the remaining soluble portion of the complex was concentrated and then purified by chromatography on a Superdex200 16/60 column followed by Resource-Q anion-exchange chromatography (GE Healthcare), as described previously (18).

### Thermostability Measurements Using Circular Dichroism Spectroscopy

The thermostabilities of Aime-128 with nine peptides were tested by CD spectroscopy. CD spectra were measured at 20°C on a Jasco J-810 spectropolarimeter equipped with a water-circulating cell holder. The protein concentration was 0.1 mg/mL in pH 8.0 Tris buffer (20 mM Tris and 50 mM NaCl). Thermal denaturation curves were determined by monitoring the CD value at 218 nm using a 1-mm optical-path-length cell as the temperature was raised from 25 to 80°C at a rate of 1°C/min. The temperature of the sample solution was directly measured with a thermistor. The fraction of unfolded protein was calculated from the mean residue ellipticity (θ) using a standard method. The unfolded fraction (%) is expressed as (θ−θ_N_)/(θ_U_−θ_N_), where θ_N_ and θ_U_ are the mean residue ellipticity values in the fully folded and fully unfolded states, respectively. The midpoint transition temperature (T_m_) was determined by fitting the data to the denaturation curves using the Origin 8.0 program (OriginLab) as described previously (39). Based on the CD spectroscopy results, the nonapeptide binding motifs of Aime-128 were extrapolated from potential CTL epitopes of viruses detected from giant panda (36).

### Crystallization and Data Collection

The purified pAime-128 complexes were ultimately concentrated to 7.5 mg/ml in crystallization buffer (20 mM Tris–HCl (pH 8.0) and 50 mM NaCl), mixed with reservoir buffer at a 1:1 ratio, and then crystallized by the hanging-drop vapor diffusion technique at 4°C. Index Kits (Hampton Research, Riverside, CA) were used to screen the crystals. After several days, pAime-128 crystals were obtained with Index solution 55 (30% (w/v) polyethylene glycol 3350, 0.05 M magnesium chloride hexahydrate and 0.1 M HEPES (pH 7.5)). Diffraction data were collected using an in-house X-ray source (Rigaku MicroMax007 desktop rotating anode X-ray generator with a Cu target operated at 40 kV and 30 mA) and an R-Axis IV imaging-plate detector at a wavelength of 0.97892 Å. In each case, the crystal was first soaked in reservoir solution containing 25% glycerol as a cryoprotectant for several seconds and then flash-cooled in a stream of gaseous nitrogen at 100 K (40). The collected intensities were indexed, integrated, corrected for absorption, scaled, and merged using the HKL2000 package (41).

### Structure Determination and Refinement

The structure of pAime-128 was solved by molecular replacement using the MOLREP program with HLA-B*5101 (PDB code, 1E27) as the search model. Extensive model building was performed by hand using COOT (42), and restrained refinement was performed using REFMAC5. Further rounds of refinement were performed using the phenix refine program implemented in the PHENIX package (43) with isotropic ADP refinement and bulk solvent modeling. The stereochemical quality of the final model was assessed with the PROCHECK program (44).The data collection and refinement statistics are listed in **Table 2**.

### Structural Analysis and Generation of Illustrations

Peptide-contacting residues were identified using the program CONTACT and were defined as residues containing an atom within 3.3 Å of the target partner. Structural illustrations and electron density-related Figs were generated using the PyMOL molecular graphics system (http://www.pymol.org). Solvent-accessible surface areas and B factors were calculated with CCP4. The multiple sequence alignment was performed by Clustal Omega (45) (https://www.ebi.ac.uk/Tools/msa/clustalo/) and ESPript 3.0 (46) (http://espript.ibcp.fr/ESPript/ESPript/). Accessible surface area (ASA) and buried surface area (BSA) were calculated by the online website PDBePISA.

### Protein Structure Accession Numbers

The crystal structures have been deposited in the Protein Data Bank (http://www.pdb.org/pdb/home/home.do) with accession numbers (5ZE5).

## ACKNOWLEDGMENTS

We acknowledge the assistance of the staff at the Shanghai Synchrotron Radiation Facility of China (SSRF).

## AUTHOR CONTRIBUTIONS

H.Y. performed the experiments; H.Y., L.M. and L.Z. analyzed the data and wrote the paper; X.L. provided constructive suggestions; and C.X. designed and supervised the study. All authors critically revised the manuscript and gave their final approval of the version to be submitted.

## SUPPLEMENTARY MATERIAL

**Supplementary Fig 1** Refolding efficiency of pAime-128 complex. Aime-128 and Aime-*β*_2_m were corefolded with different nonapeptides. The refolded products were purified by gel filtration chromatography and anion-exchange chromatography. pAime-128 complex b curves are shown in blue. The insets show reducing SDS-PAGE gels (15%) of the peaks that are labeled on the curve. Lane M contains molecular mass markers (labeled in kD). (**A**) Refolding efficiency of Aime-128 and Aime-*β*_2_m with NGY9. (**B**) Refolding efficiency of Aime-128 and Aime-*β*_2_m with TSV9. (**C**) Refolding efficiency of Aime-128 and Aime-*β*_2_m with NTY9. (**D**) Refolding efficiency of Aime-128 and Aime-*β*_2_m with EVW9.

